# Leveraging biological insight and environmental variation to improve phenotypic prediction: Integrating crop growth models (CGM) with whole genome prediction (WGP)

**DOI:** 10.1101/100057

**Authors:** C.D Messina, F. Technow, T. Tang, R. Totir, C. Gho, M. Cooper

**Affiliations:** DuPont Pioneer, 8503 NW 62nd Avenue, Johnston, IA 50131, USA.; DuPont Pioneer, 596779 County Road 59N, Woodstock, Ontario N4S 7W1, Canada.; DuPont Pioneer, Semillas Pioneer Chile Ltda, Santa Filomena 1609 – Buin, PO Box 267, Chile.

**Keywords:** whole genome prediction, crop growth model, maize, breeding, physiology, genotype by environment interactions, phenomics, phenotyping

## Abstract

A successful strategy for prediction of crop yield that accounts for the effects of genotype and environment will open up many opportunities for enhancing the productivity of agricultural systems. Crop growth models (CGMs) have a history of application for crop management decision support. Recently whole genome prediction (WGP) methodologies have been developed and applied in breeding to enable prediction of crop traits for new genotypes and thus increase the size of plant breeding programs without the need to expand expensive field testing. The presence of Genotype-by-Environment-by-Management (G×E×M) interactions for yield presents a significant challenge for the development of prediction technologies for both product development by breeding and product placement within agricultural production systems. The integration of a CGM into the algorithm for whole genome prediction WGP, referred to as CGM-WGP, has opened up the potential for prediction of G×E×M interactions for breeding and product placement applications. Here a combination of simulation and empirical studies are used to explain how the CGM-WGP methodology works and to demonstrate successful reduction to practice for applications to maize breeding and product placement recommendation in the US corn belt.

## 1. Introduction

Whole Genome Prediction (WGP) is a set of quantitative genetic methodologies that enables prediction of the breeding value of an individual, created from one or more reference populations, based on its genetic makeup and pedigree relations. In combination with high throughput genotyping and phenotyping WGP has brought unprecedented change to the scale of plant breeding (Heffner et al., 2009; Lorenz et al., 2011; Cooper et al., 2014). Unlike the near past, today for most large commercial breeding programs germplasm is evaluated using WGP and only a fraction of the new hybrids that can be created are evaluated in multi-environment trials (MET, Heffner et al., 2009; Cooper et al., 2014). WGP enabled breeders to increase the effective size of their breeding programs without the need to increase the scale of field phenotyping and by doing so accelerated the rate of genetic gain.

The WGP methodology seeks to simultaneously estimate the allelic values at all available polymorphic marker loci across the genome (Meuwissen et al., 2001). Bayesian approaches seek to estimate the posterior distribution of marker effects by means of calculation of a likelihood function and a prior distribution of such effects. This is a data driven process that leverages the availability of large datasets routinely generated in commercial breeding programs. Datasets often comprise multiple years and large samples of genotypes from reference populations with the expectation that these are a representative sample of the germplasm and the target population of environments (TPE, Cooper et al., 2014). The mechanics of the method separates individuals within populations under selection into the WGP estimation set, individuals in this set have both phenotypic and genotypic information, and the prediction set, individuals in this set only have genotypic information (see Heffner et al., 2009, Lorenz et al., 2011). Various statistical methodologies were developed to enable WGP, among which selection of a suitable method depends on genetic architecture of the trait, the ability to demonstrate and model non-linear interactions and the population structure (Lorenz et al., 2011). Recently, the use of biological frameworks to model non-linear genetic effects was advocated to improve prediction accuracy (Marjoram et al., 2014, Technow et al., 2015). A necessary condition to leverage such biological insight in WGP is that that knowledge be encapsulated in the form of quantitative functions that transparently map markers to biological function (Messina et al., 2011; Cooper et al., 2005) and be linked to prediction algorithms (Technow et al., 2015) to generate the necessary boundary conditions to accurately execute the biological algorithms (Cooper et al., 2016).

Crop Growth Models (CGMs), structured on principles of resource capture (e.g., solar radiation, water, nitrogen), use efficiency and allocation to organs of economic value, provide the biological framework for phenotypic prediction of relevant quantitative traits for breeding (Cooper et al., 2009). A general mathematical framework to develop models for phenotypic prediction with genetic information based on the E(N:K) family of models (Cooper et al., 2005) has been developed (Messina et al., 2011). Examples of implementations include models of leaf elongation rate and flowering time in maize (Welcker et al., 2007; Dong et al., 2012), and oat and soybean growth and development (Yin et al., 2004; Messina et al., 2006). But the parameterization of CGMs at a scale that is required to support breeding programs using these approaches proved challenging because of the difficulty of phenotyping important physiological traits (Messina et al., 2011). This phenotyping bottleneck currently limits the applicability of CGM to augment plant breeding. Advances in phenomics will undoubtedly generate very large datasets (Furbank and Tester, 2011; Fahlgren et al., 2015) bringing opportunity to improve models, and challenges to the utilization of these sources of information. Efficient analytical methods that reduce the phenotyping requirements and directly integrate biological models, such as CGMs, into the prediction algorithms in a single step, are naturally well positioned to fully take advantage of improvements in phenomics.

The CGM-WGP methodology uses a CGM as part of the calculation of the likelihood function step in an otherwise standard WGP algorithm (Technow et al., 2015). The relationship between marker effects and yield is established through the estimation of marker effects and biological parameters within the CGM. Because the selected genetic models result in the modulation of the strength of the relationship between the environment and a physiological process, and/or different physiological processes, genotype-by-environment (G×E), epistasis in the form of trait-by-trait genotype-by-genotype (G×G) and genotype-by-environment-by-management (G×E×M) interactions are all emergent properties from the genetic variation effects in the functional equations that relate traits and the environment (Messina et al., 2011; Cooper et al., 2009). Technow et al. (2015) demonstrated this concept using a simulation experiment, while Cooper et al. (2016) demonstrated the feasibility of implementing CGM-WGP within a functional breeding program. However, a demonstration that CGM-WGP prediction accuracy is higher than benchmark methodologies in at least one realistic breeding case study is lacking. It was hypothesized that the difference in predictive accuracy between CGM-WGP and benchmark WGP algorithms will increase with an increasing role of G×E interaction in determination of performance, and that parameterization of the CGM-WGP models will improve with the contrast between environmental conditions (Cooper et al., 2016).

Because training of CGM-WGP involves the estimation of biological parameters that regulate physiological behavior, the CGM-WGP not only can output predictions for yield for the set of environments and hybrids used for training as for other prediction methods such as BayesA, but also enables the breeder to exercise the CGM to make predictions to evaluate hybrids that have never been empirically evaluated in the field at a large scale, through computer simulation. This simulation step enables extending predictions from a few G×M or G×E cases intrinsic to the training sets to virtually the target population of environments, here considered broadly to include agronomic practices.

The objectives of this paper are to: 1) extend CGM-WGP methodology to train models using data from multiple environments, 2) evaluate, using both synthetic and experimental data from a maize drought breeding program, whether CGM-WGP methodology can enable improved phenotypic prediction when G×E interactions are an important determinant of performance, 3) demonstrate virtual breeding by means of use of CGM-WGP and computer simulation, and 4) facilitate the dialogue between breeders, crop physiologists and modelers.

## 2. Materials and Methods

### 2.1 Hierarchical model

The model fitted to the data (Fig. 1) uses the likelihood derived from the generalized E(N:K) model (Cooper et al., 2005; Messina et al. 2011) as,

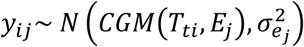

Where *y_ij_* is the yield of individual *i* in environment *j*, and *T_ti_* is the unobserved value for physiological trait *t* (e.g., AMAX; a measure of maize canopy size based on the area of the largest leaf) for individual *i*. The environmental inputs (e.g., soil type, temperature) of environment *j* are represented by *E_j_*, and 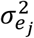 denotes the residual variance for yield in that environment. Finally *N* denotes the Gaussian density function and *CGM* the crop growth model. Thus, *CGM*(*T_ti_*, *E_j_*) denotes the simulated yield of individual *i* in environment *j*, as determined by the states of the physiological traits *t* for individual *i*, *T_ti_*.

**Figure 1.**
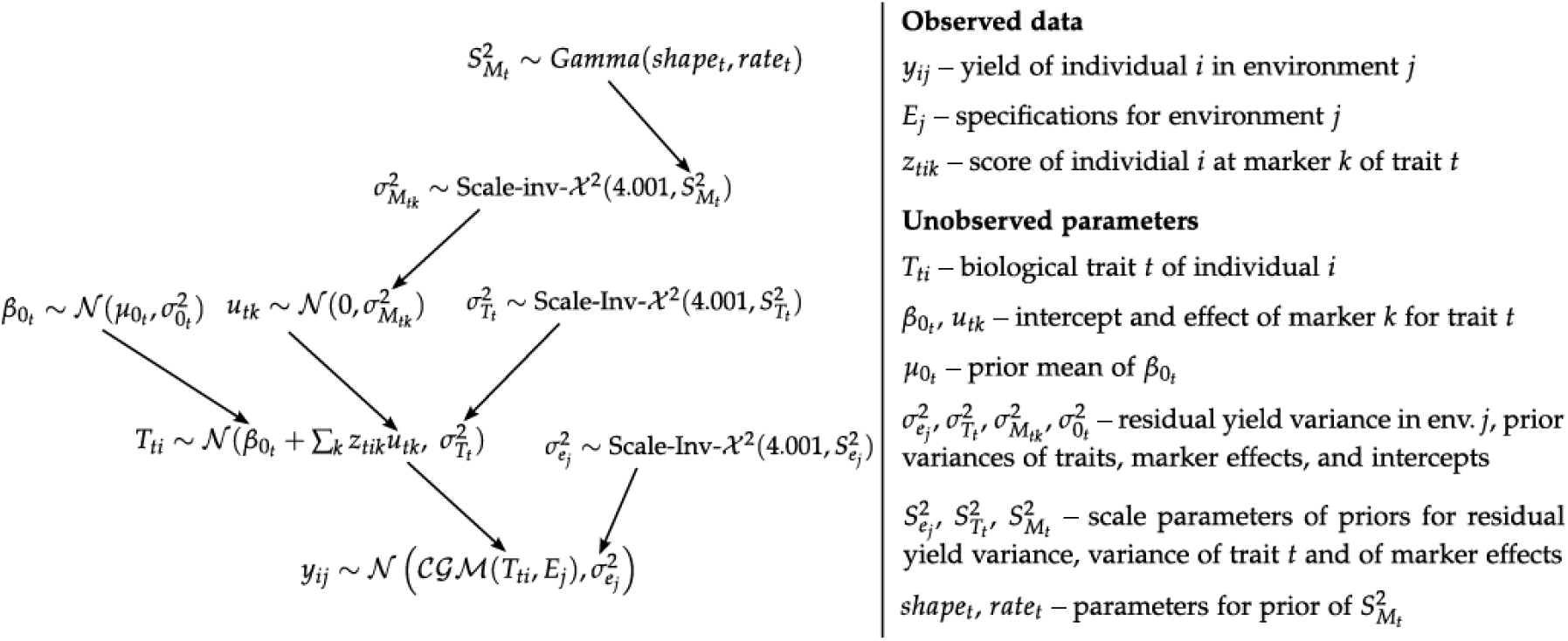
Hierarchical crop growth model (CGM) whole genome prediction (WGP) algorithm.

The prior for the unobserved *T_ti_* was

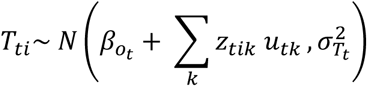

where *β_o_t__* is a trait specific intercept, *z_tik_* denotes the marker score of individual *i* at marker *k* for trait *t* and *u_tk_* the effect of marker *k* for trait *t*. The trait specific prior variance was 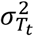.

The prior distributions for the marker effects correspond to those from the widely used whole genome prediction model ‘BayesA’ (Meuwissen et al., 2001). Briefly, the marker effects *u_tk_* were associated with a Normal prior distribution with mean 0 and marker specific variancea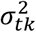, which itself was associated with scaled inverse Chi-square prior distribution with 4.1 degrees of freedom and trait specific scaling factor 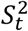. The prior distribution for the parameter 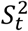 was a Gamma distribution with constant parameters (Yang and Tempelman, 2012). The prior for the trait specific intercept *β_o_t__* was a Normal distribution with mean *μ*_0_*t*__ and variance 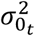, both of which were constants. The prior of 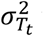 was again a scaled inverse Chi square distribution with 4.001 degrees of freedom and constant scaling factor 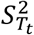. Finally, the prior of the environment specific residual variance 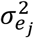 was also a scaled inverse chi-square distribution with 4.001 degrees of freedom and constant scaling factor 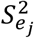. Thus, the CGM-WGP model can be understood as a generalized linear model version of the whole genome prediction model ‘BayesA’ (Meuwissen et al., 2001), in which the crop growth model acts as the link function.

The constants were derived from rough prior estimates of the mean (*m_t_*) and variance (*v_t_*) of physiological trait *t* within the germplasm under consideration. From these, the scaling factor of the prior of the physiological traits was calculated as 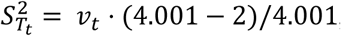, which results in a scaled inverse chi-square prior distribution with mean *v_t_*. The prior parameters for the trait specific intercepts *β_0_t__* were *μ*_0_*t*__ = *m_t_* and 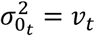. The parameters of the Gamma prior distribution of 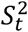, the scaling factor of the marker specific variances, were calculated as follows. First the variance of the additive effect of a random marker locus was calculated according to Habier et al. (2011) as 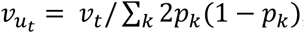, where *P_k_* is the allele frequency of marker *k*. From this we calculated the prior expected value of 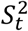 as 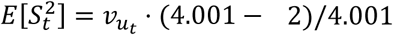, the values of *shape_t_* and *rate_t_* were then chosen in such a way that the mean of the Gamma prior distribution was 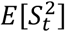 and its variance 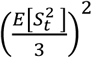.

### 2.2 Metropolis-within-Gibbs sampling algorithm

A ‘Metropolis-Hastings within Gibbs’ algorithm was implemented to sample from the posterior distribution of all parameters (Gelman et al., 2004; see Wallach et al., (2012) for an application to estimating crop growth model parameters). Briefly, the Gibbs sampler (Gelman et al., 2004) is a Markov chain algorithm for high-dimensional parameter spaces. Because the algorithm samples parameters sequentially from the conditional (on the data and all other parameters) posterior distribution of each, the algorithm is highly efficient. With the exception of *T_ti_*, the conditional posterior distributions of all parameters in CGM-WGP are recognizable distributions that can be sampled from directly. A Metropolis-Hastings step, which is an accept-reject algorithm that can sample from any distribution, was therefore included in the final algorithm to sample *T_ti_*. The Gibbs sampler use in this study is identical to the Gibbs algorithm generally used for BayesA models (Meuwissen et al., 2001; Yang and Tempelman, 2012), except for the sampling of *T_ti_* with the Metropolis-Hastings algorithm.

Conditional on the marker effects *u_tk_*, the physiological traits *T_ti_* of one individual are independent of those of the others and can hence be sampled sequentially. The different traits for each individual, however, are not and have to be sampled jointly. This was done as described by Wallach et al. (2012), with the exception that a change of variable was implemented to sample the parameters in the space of the natural logarithm. This is a common technique when the parameter values in the original space have to be positive (Gelman et al., 2004). The Metropolis-Hastings step was run for two subsequent iterations to improve convergence.

Gibbs sampling chains were run for 200 thousand iterations. The first half of each chain was discarded as ‘burn-in’ and samples from every 100^th^ iteration thereafter were stored, thus resulting in 1000 stored samples. Convergence was monitored by inspecting graphical diagnostics such as implemented in the R package ‘coda’ (Plummer et al., 2006). Increasing the chain length did not seem to improve prediction accuracy. The algorithm was implemented as a C routine and embedded in the R statistical software environment (R Core Team, 2014).

### 2.3 Prediction of yield from CGM-WGP results

The predicted yield of an untested individual *i*′ in environment *j*′ was obtained from the posterior samples of the marker effects *u_tk_* and the intercepts *β_0_t__*. From each posterior sample the values of the physiological traits were calculated as 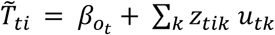. Those values were then entered into the CGM together with the inputs of the environment, resulting in one simulated yield value per posterior sample. Those samples represent the posterior predictive distribution of yield for individual *i*′ in environment *j*′. The mean of this distribution was used as the predicted value. The algorithm BayesA (Meuwissen et al., 2001) was utilized as a reference method that predicts yield using marker information only. i.e., 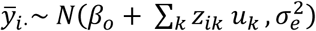, with *y̅_i_*. denoting the yield of individual *i* averaged over all environments considered for estimation. The prior of the marker effects *u_k_* was 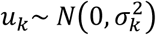 and standard, uninformative prior distributions were used for 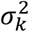 and 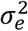. The BayesA Gibbs-sampler was run for 50000 iterations, of which the first 25000 were discarded and samples from only every 25^th^ subsequent iteration stored. In this study, BayesA served as a purely statistical reference method relative to which we measure the benefit of modeling G×E interactions with CGM-WGP.

### 2.4 Crop growth model

The CGM connected to the WGP algorithm is a mechanistic model based on the concept of radiation and water capture and use efficiencies, and mass allocation to reproductive growth (Muchow et al., 1990; Muchow and Sinclair, 1991). Crop and canopy development are simulated as a function of thermal time (TT, °C) with base temperature for preflowering equal to 8°C and postflowering equal to 0°C (Muchow et al., 1990). Growth (W) is calculated as the product of light interception (LI) and radiation use efficiency (RUE), W=LI × RUE. The value for RUE was set to 1.85 g MJ^−1^ after Hammer et al. (2009), unless explicitly changed for specific studies described below. Light interception is estimated from the leaf area per plant (LP), plant density (PD) and canopy attributes described by the coefficient of extinction, which has set to 0.4, LI=1-exp(-0.4 × LP × PD). Leaf area per plant is modeled solely as a function of the size of the largest leaf (AMAX, Birch et al., 1998); that is, other parameters are kept constant as in Muchow et al. (1990). Transpiration (TR) is calculated as a function of W, vapor pressure deficit (VPD), transpiration efficiency coefficient (TE=9 kPa, Tanner and Sinclair, 1983), and the expression of the limited transpiration trait (Sinclair et al., 2005; Messina et al., 2015) on an hourly (h) time step,

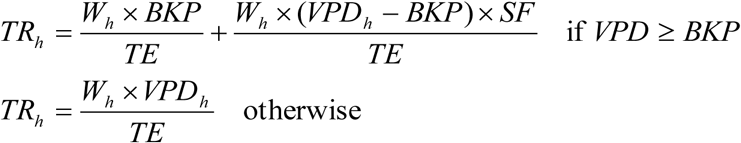

where BKP is the breakpoint and SF is the sensitivity of TR to VPD. The parameter SF was set at 0.3 in order to incorporate relevant functional biological behavior (Gholipoor et al., 2013; Choudhary et al., 2014; Shekoofa et al., 2015). When hourly VPD exceeds BKP, hourly growth W_h_ is updated to conform to hourly TR and a constant TE,

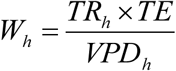

Water deficit effect on growth is modeled through the coefficient water supply-to-demand (SD) ratio. Water demand is calculated as TR in the absence of soil water deficit. Water supply is estimated as the sum across all soil layers of soil water content minus the soil water content at the lower limit times the *k_l_* as in Robertson et al. (1993), where the *k_l_* coefficient was set at 0.08 (Dardanelli et al., 1997; Hammer et al., 2009). Soil water content is estimated using a multilayer model. Grain yield is simulated as the linear increase of harvest index (HI) during seed fill. Reduction in postflowering growth due to stress at flowering time is simulated by modeling the attainable HI from silk numbers exerted at the onset of the increase in HI, as defined in Muchow and Sinclair (1990), and the maximum silk numbers (SNM), a value that can vary among genotypes. Silk number (SN) is estimated from the ear mass (EM) as, SN = SNM × (1 - exp(-0.14 × (EM – MEB)), where MEB is a parameter characteristic of genotype (Cooper et al., 2014), and EM is modeled using an exponential function of TT and a stress factor directly proportional to SD. The parameter MEB corresponds to the threshold of ear biomass below which silk elongation is not fast enough to exert silks beyond the ear husk. Chenu et al. (2009) used a similar approximation that included a threshold similar to MEB to model the connection between QTL controlling silk elongation, anthesis-silking interval, which is a crop attribute closely related to yield under water limited conditions (Bolaños and Edmeades, 1993; Cooper et al., 2014), and yield. For further discussion of CGMs Soltani and Sinclair (2012) provide an extensive treatment of simple mechanistic crop models.

### 2.5 Multi-environment trial simulation

A multi-environment trial simulation experiment based on three environments was created to assess the CGM-WGP methodology. To characterize the three environments and quantify water availability during the crop cycle, water SD ratios were calculated on a daily time step following the procedures described by Cooper et al. (2016). Consequently it was determined that the simulated multi-environment trial comprised of three environments with different degrees of water deficit occurring in Johnston, Iowa, US (41.684 °N, −93.508°W) in 1988, 2012 and 2010, herein referred to as water limited 1 (WL1), water limited 2 (WL2) and NWL (not water limited), respectively. The soil depth, soil water holding capacity, and *k_l_* were 2.2 m, 0.13 cm^3^.cm^−3^, and 0.08, respectively. Planting dates were 4/29/1988, 4/28/2010, and 5/4/2012. Plant population was set at 6.5 pl m^−2^. Daily meteorological records were from the National Oceanic and Atmospheric Administration (NOAA; Bell et al., 2013). Genotypic variation for a set of physiological traits was considered for a Doubled Haploid (DH) population of lines. The DH lines were evaluated as F1 hybrids in combination with an inbred from a complementary heterotic group. Parameters describing these traits for an individual DH within a population were allowed to vary within the following intervals: [700 < AMAX< 1100 cm^2^] (Elings, 2000), [1.6 < RUE<2.1 g MJ^−1^] (Sinclair and Muchow, 1999), [0.5<MEB<1.0 g] (Cooper et al., 2014), and [1.5<BKP<2.5 kPa] (Messina et al., 2015).

The DH genotypes of the DH lines were generated *in silico* in two step process, similar to the one used by Technow et al. (2014a) for simulating a DH maize breeding population. In step one, an ancestral population of 50 inbred lines was stochastically simulated. The genome consisted of 10 chromosomes with lengths between 0.75 and 1.25 Morgan (M). Two hundred evenly spaced biallelic marker loci were placed on each chromosome. The total number of markers was thus 2000. An additional 40 loci were randomly distributed across all chromosomes that served the purpose of simulating quantitative trait loci (QTL). Historical linkage disequilibrium (LD) between markers as well as their allele frequencies was simulated with the method described by Montana (2005). The simulated expected LD (measured as squared correlation) between markers followed an exponential decay, such that it halved approximately every 0.1 Morgan. For two marker loci *t* Morgan apart, the expected LD followed 0.5 · 2^(–*t*/0.1)^ + 0.1 · 2^(–*t*/0.5)^. Exponential decay curves mirror the LD decay curves observed in maize (Technow et al., 2014b). Minor allele frequencies were drawn at random from the (0,0.5) interval. This population was then random mated for three generations to generate a pedigree and sub-population structure. The population size was thereby kept constant at 50. After the last generation 1000 doubled haploid lines were generated through meiosis followed by a chromosome doubling step.

The values for the physiological traits were simulated according to Technow et al. (2015). A unique set of ten of the 40 loci set aside as QTL were assigned to each of the four physiological traits: AMAX, RUE, MEB, and BKP. The additive substitution effect of each QTL was drawn from a Standard Normal distribution and raw genetic scores calculated for each trait by summing the effect of the QTL according to the genotypes of the DH lines. These scores were then rescaled linearly to the ranges described above.

The true grain yield value of a DH line *i* in environment *j* was then simulated by executing the CGM with the physiological trait values of the DH and the appropriate environmental inputs. For those DH that became part of the estimation set we also simulated observed phenotypic yield values by adding a Normal noise variable to the true values. The variance of this variable was chosen in such a way that the within environment heritability was equal to 0.66.

### 2.6 Whole genome prediction application

Estimation sets for training the CGM-WGP model were constructed from one, two or all three environments. For the single location estimation sets (WL1, WL2 or NWL) the phenotypic yield data of a random sample of 500 of the DH lines in that environment were used. For two and three environment estimation sets, yield data from 250 DH lines in the two environments case and 166 DH lines in the three environments case were used. The total number of data points was thus equal in all cases. Values for 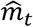 and 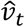, which are required for calculation of constants and starting values, were computed from estimates obtained from a random sample of 25 DH lines, which were assumed to be phenotyped for the traits but were not used as part of the estimation set. The CGM-WGP algorithm was then run as described and the marker effects used to predict the yield performance of all DH lines within the validation set in all three environments, also as described. Prediction accuracy was then calculated as the Pearson correlation between predicted and true yield in each environment. Because the true values of the physiological traits were known, it was possible to assess their prediction accuracy by calculating the correlation between their predicted and true values. BayesA methodology was utilized as a reference method and it is described above. The whole process, including the data simulation, was repeated 75 times for each environment combination. The resulting distribution of the accuracy statistics was summarized by the mean and standard deviation.

### 2.7 Empirical breeding experiment

For the empirical evaluation the CGM-WGP and BayesA methods prediction accuracies were compared using four DH populations that were tested in an experiment that comprised of a non-water-limited (NWL) environment and a water-limited (WL) environment. Both field environments were created at the DuPont Pioneer Viluco research station, which is located in Chile (-33.797 °S, −70.807 °W), in the 2012/13 season. The contrasting water environments were created by controlled application of quantity and timing of water during the crop cycle. As for the simulation experiment, water SD ratios were estimated on a daily time step during the crop cycle for the NWL and WL environments to quantify the impact of the different irrigation regimes that were used to create the contrasting water environments.

The four DH populations, referred to as DHPop1, DHPop2, DHPop3, DHPop4, were created from biparental crosses between six inbred lines, referred to as I1, I2,…, I6. The two parents of DHPop1, I1 and I2, were different from the four inbred lines, I3, I4, I5 and I6, used as parents to create the other three DH populations. One of the four remaining inbred lines, I3, was used as a common parent for all three remaining DH populations; DHPop2 parents I3/I4, DHPop3 parents I3/I5 and DHPop4 parents I3/I6. Thus, there is a closer pedigree relationship, based on the half-sib structure, between DHPop2, DHPop3 and DHPop4 than there is between any of these three populations and DHPop1. Each of the four populations was represented in the experiment by 76 to 105 DH lines. The DH lines were each genotyped with approximately 1600 polymorphic Single Nucleotide Polymorphism (SNP) markers.

For the experimental evaluation of grain yield, testcross hybrids were created for all of the DH lines using a common inbred tester line that was selected from a complementary heterotic group. Thus, while for discussion purposes we refer to DH lines they were evaluated for grain yield as hybrids in the NWL and WL environments. Grain yield was measured for all DH lines from two-row plots. The two-row plots were 4.5m long with 0.75m spacing between rows. The number of plants within the rows was managed to represent a plant population of 7.9 pl. m^−2^. To manage irrigation quantity and timing drip tape was installed at a depth of approximately 0.1m and approximately 0.1m to the side of each plot row at the time of planting of the experiment. At maturity the total grain yield of a plot was measured by harvesting all ears from within the plot using a two-plot combine harvester. The total weight and moisture content of the grain from a plot were measured at the time of harvest and the harvested plot weight was converted to grain yield on an area basis at 15% moisture.

The experimental design in each environment was a row-column design with two replicates. The grain yield data were analyzed using the ASREML mixed model software (Gilmour et al. 2009). For the mixed model analysis the two environments, NWL and WL, were treated as fixed effects and the DH lines were treated as a random sample of lines from within their respective population. To investigate the magnitude of G×E interaction for grain yield between the NWL and WL environments for each of the four DH populations a combined analysis of variance was conducted following the recommendations of van Eeuwijk et al. (2001). When significant G×E interactions were detected the genetic correlation was estimated for grain yield between the two environments (Falconer and Mackay 1996). For reference, the genetic correlation can range from 1.0 to −1.0. A genetic correlation of 1.0 indicates there were no G×E interactions. Conversely, a genetic correlation of −1.0 indicates strong G×E interactions with complete rank reversal of the DH lines between the two environments. For the selected mixed model analysis of variance Best Linear Unbiased Predictors (BLUPs) were computed for grain yield of each of the DH lines in the NWL and WL environments. These grain yield BLUPs were then used for the CGM-WGP and BayesA prediction analyses.

A leave-one-family-out cross-validation was conducted to assess prediction accuracy for grain yield. Here the CGM-WGP model was selected using an estimation data set based on the yield data from both the NWL and WL environments for three of the four DH populations and then used to predict the yield values of the fourth DH population for both the NWL and WL environments. This process was repeated until each population was left out once. Since the estimation set comprised yield data from both the yield in the NWL and WL environments this is a multi-environment estimation set. Prediction accuracy for the lines of the DH population left out of the estimation set was calculated as the correlation between predicted and observed yield BLUP values separately for both the NWL and WL environments.

The physiological traits for the empirical study were the same as those used for the simulation study; AMAX, RUE, MEB and BKP. Prior estimates of physiological trait means and variances 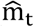 and 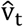 covered the typically observed biological ranges. The values of the means were 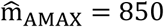, 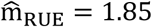, 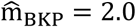, and 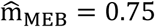 and those of the variances 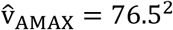, 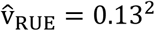, 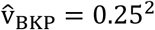, and 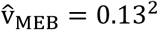. The values of 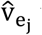 were 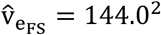 and 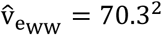. Both of which were obtained from an ASREML analysis of the original data.

The classical BayesA WGP model (Meuwissen et al. 2001) was used as a reference method relative to which we measure the benefit of modeling G×E interactions with CGM-WGP. The BayesA model was applied directly to the BLUP yield average over the two environments.

### 2.8 Large scale evaluation of hybrids never tested in the field

The CGM-WGP methodology can be used to make predictions within the set of experiments used for model training and evaluation, as demonstrated by Cooper et al. (2016). This application is comparable with BayesA and other statistical based methodologies (Heffner et al. 2009, Lorenz et al., 2011). Because CGM-WGP estimates the value of the alleles for each polymorphic marker locus for each physiological parameter included for CGM-WGP model training, it is possible to generate predictions for any combination of management and environment for any individual that has been genotyped and belongs to the genetic inference space. A simulation experiment using genetic parameters estimated for individuals from a breeding population (see above) and environments included in the TPE (Messina et al., 2015) was conducted to demonstrate the feasibility of augmenting field evaluation with in-silico evaluation of untested genotypes at large scale. Parameters to run the mechanistic CGM model were estimated for individuals from a breeding population using the method and breeding experiments described above. Environment and management inputs to run the CGM in 2263 30 x 30 km grids within the maize growing region for 1988 and 2014 are described in Messina et al. (2015). Briefly, weather data were from NOAA. Solar radiation was estimated from temperature records (Bristow and Campbell, 1984) with parameters provided by Mud Springs Geographers, Inc, and hourly vapor pressure deficit (VPD) was estimated from daily temperature assuming an harmonic change in daily cycle (Monteith and Unsworth, 1990). Soil depth was from the STATSGO database (United States Department of Agriculture, 2015) and used to estimate total soil water holding capacity by multiplying it by a constant (0.13 cm^−3^ cm^−3^) volumetric fraction of available soil water. Crop management data that includes plant population, planting date and maturity group were from DuPont Pioneer data bases.

## 3. Results and Discussion

### 3.1 *Physiological determinants of* G×E *interactions for yield*

The simulation study included three environments contrasting for water availability, as characterized by the water supply to demand ratio index (Fig. 2). Water deficit was largest at flowering time in WL1 and at grain filling in WL2. Simulations for the year 2010 (NWL) indicated absence of water stress with SD equal to 1 throughout the growing season (SD for NWL not shown on Fig. 2). This combination of environments set the conditions to observe differential effects of physiological traits on yield (Fig. 3) and consequently create a case study to test the CGM-WGP and BayesA algorithms for ability to construct predictive genetic models (Tables 1, 2).

**Figure 2.**
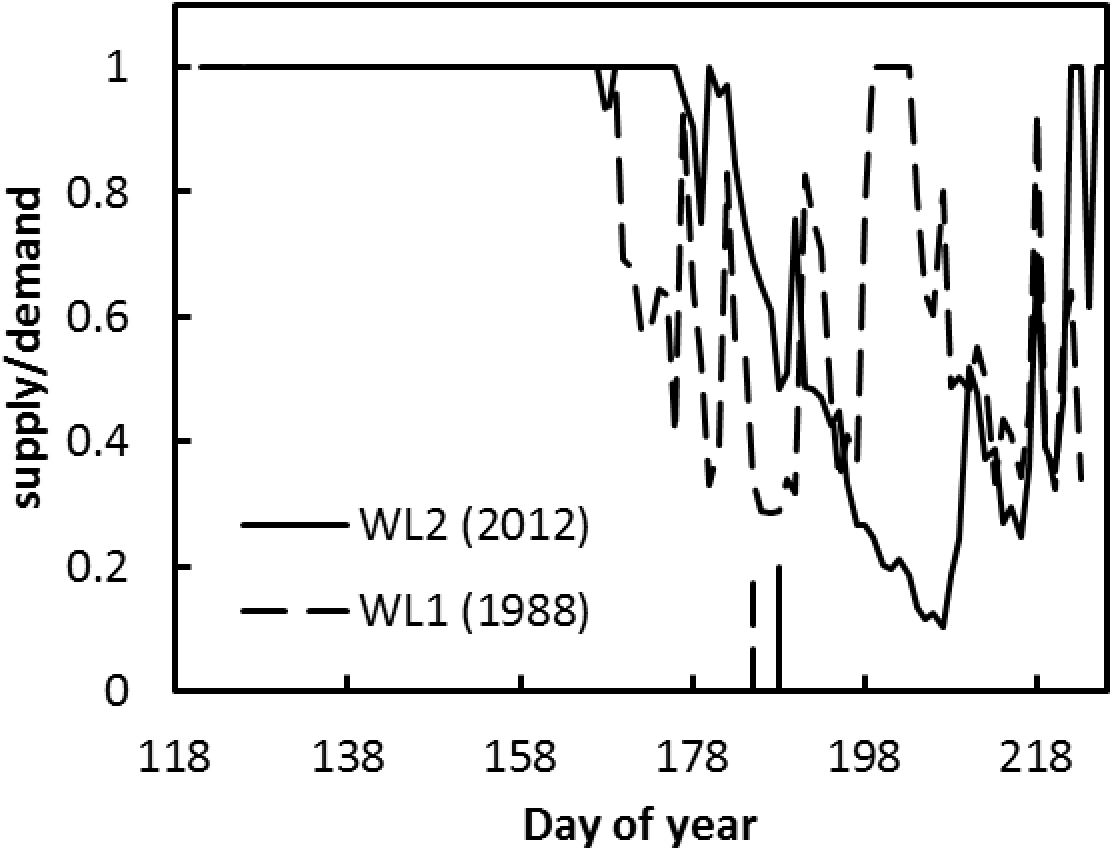
Water supply to demand ratios estimated on a daily time step for Johnston, IA calculated for 1988 (water limited, WL1) and 2012 (water limited 2, WL2) using a crop growth model as a function of day of year. Simulated shedding dates for 1988 and 2012 indicated as vertical bars. Model parameters were set to means of values utilized in sensitivity analyses, size of the largest leaf in the leaf area profile (AMAX)=950, ear size at silking (MEB)=0.8, radiation use efficiency (RUE)=1.8, and limited transpiration breakpoint (BKP)=2.25.

**Figure 3.**
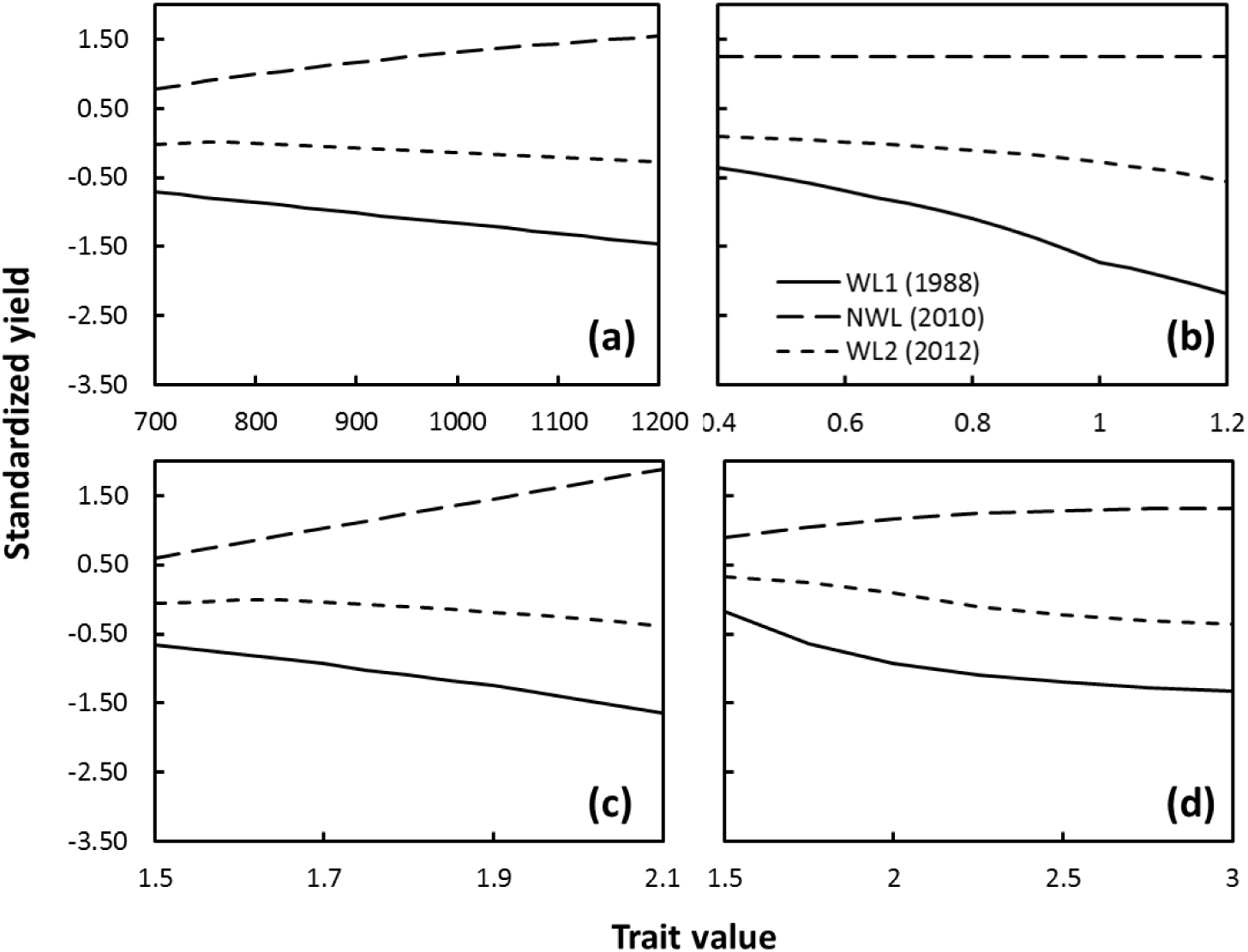
Response of standardized yield to variation in crop growth model parameters: (a) size of the largest leaf in the leaf area profile (AMAX, cm^2^), (b) ear size at silking (MEB, g), (c) radiation use efficiency (RUE, g MJ^−1^), and (d) limited transpiration breakpoint (BKP, kPa) for water limited environments (WL1) 1988 and (WL2) 2010, and not water limited environment (NWL) 2012 at Johnston, IA USA (See Figure 2 for dynamics in supply/demand ratio). Yields were standardized relative to the overall mean (1054 g m^−2^) and standard deviation (407 g m^−2^).

**Table 1.** Mean prediction accuracies (± standard deviation) for yield estimated by the correlation coefficient *r* for CGM-WGP prediction methodology and the reference method BayesA.

| Estimation<br>environment | Prediction environment |  |  |  |  |  |
| --- | --- | --- | --- | --- | --- | --- |
|  | CGM-WGP |  |  | BayesA |  |  |
|  | WL1 | WL2 | NWL | WL1 | WL2 | NWL |
| WL1(1988) | 0.85 ±0.03 | 0.51±0.13 | 0.60±0.13 | 0.84 ±0.03 | 0.38±0.17 | -0.62±0.14 |
| WL2(2012) | 0.34±0.43 | 0.79±0.06 | 0.23±0.42 | 0.44±0.14 | 0.78±0.06 | 0.06±0.28 |
| NWL(2010) | 0.57±0.11 | 0.22±0.16 | 0.88±0.02 | -0.60±0.11 | 0.01±0.26 | 0.88±0.02 |

**Table 2.** Mean prediction accuracies (± standard deviation) for yield estimated by the correlation coefficient *r* for CGM-WGP prediction methodology and the reference methodBayesA and nine cases defined by a unique combination of two or three environments utilized to train the algorithms and one environment of utilized for evaluation of accuracy. Environments include two water limited (WL) and one not water limited (NWL) conditions.

| Estimation<br>environment | Prediction environment |  |  |  |  |  |
| --- | --- | --- | --- | --- | --- | --- |
|  | WL1(1988) | WL2(2012) | NWL(2010) | WL1(1988) | WL2(2012) | NWL(2010) |
|  | ----- CGM-WGP ----- |  |  | ----- BayesA ----- |  |  |
| WL1 + WL2 | 0.79±0.04 | 0.75±0.05 | 0.67±0.08 | 0.77±0.04 | 0.51±0.13 | -0.50±0.15 |
| WL2 + NWL | 0.72±0.09 | 0.75±0.06 | 0.81±0.04 | -0.39±0.16 | 0.26±0.25 | 0.75±0.07 |
| WL1 + WL2 +<br>NWL | 0.76±0.05 | 0.71±0.06 | 0.77±0.04 | 0.43±0.13 | 0.53±0.11 | -0.01±0.2 |

Figure 3 shows the conditional influence of the four physiological traits on yield in the three environments when holding the other traits constant at their mean and yield is standardized relative to the distribution when all traits are varying. Because of the occurrence of stress around flowering time (Fig. 2), MEB has the largest impact on yield in environment WL1 (Fig. 3b). Yield and drought tolerance in maize decreased in WL1 with increasing MEB, as demonstrated in previous studies (Messina et al., 2011, Cooper et al. 2014). With increasing AMAX (Fig. 3a), RUE (Fig. 3c) and BKP (Fig. 3d) traits yield increased in the NWL environment and decreased in the drought stress environments WL1 and WL2. These results are consistent with those observed in other studies (Sinclair and Muchow, 2001; Messina et al., 2011; Messina et al., 2015). The traits AMAX, RUE and BKP regulate crop canopy level transpiration thus the water use dynamics during the growing season. Water use decreases with decreasing values of any of these traits, which is a mechanism to conserve water during the vegetative period. When water deficit occurs at flowering time, water conservation may have a large impact on yield (Cooper et al. 2014a). Because of the high sensitivity of silking to water deficit (Hall et al., 1982; Bolaños and Edmeades, 1993), the greater intensity of water deficit at flowering time in WL1 (Fig. 2) yield in WL1 decreased with increasing AMAX, RUE and BKP traits more than in WL2 (Fig. 3). At a constant value of AMAX, RUE, BKP or MEB, simulated yield using the CGM can increase, decrease or show no change depending on the environment. This G×E interaction for yield, resulting from the interplay between physiological process and the environment, was previously referred as functional emergent behavior (Hammer et al., 2006). This attribute of the CGM to generate variable outputs conditional to a vector of constants that characterize physiological traits is what make these algorithms potentially useful functions to construct simple additive predictive genetic models for traits incorporated into the CGM that can produce complex outputs that reproduce G×E interactions and fitness landscapes for yield (Messina et al., 2011; Hammer et al., 2006; Hammer et al., 2010).

### 3.2 CGM-WGP incorporates biological insight within genomic prediction algorithms and increase prediction accuracy

The simulation results reported in this study introduce enhanced realism relative to Technow et al. (2015) with yield responses to physiological trait variation more subtle and linear than in the previous study (Fig. 3). The average prediction accuracy of CGM-WGP when trained in one, two or three environments was 0.55 (Table 1), 0.75 and 0.75 (Table 2), respectively. In contrast, the average prediction accuracy for BayesA was 0.24 for one (Table 1), 0.23 for two and 0.32 for three (Table 2) training environments, respectively. Despite the absence of major nonlinear yield response to variation in physiological traits, as in Technow et al. (2015), the results from this study support the conclusion that CGM-WGP increased prediction accuracy over the BayesA method by incorporating biological insight into the prediction algorithm.

Although the CGM-WGP average accuracy was greater than for BayesA, there were cases where BayesA accuracy could be equal or greater than CGM-WGP’s (Table 1), for example, when the prediction and the estimation environments were alike due to the common effects of drought (WL1, WL2). Cooper et al. (2016) reported similar results for an application of CGM-WGP to two drought environments that were part of a MET from within a breeding program. In contrast, there were no cases where the accuracy of BayesA was greater than that for CGM-WGP when the estimation environment differed from the prediction environment (e.g., WL1 vs. NWL). In the presence of the emergent G×E interactions at the yield level due to the contrasting influences of the physiological traits in the different environments (Fig. 3) BayesA accuracy could be negative (Table 1). For example, when BayesA was trained in WL1 and predictions were made in NWL, the mean prediction accuracy was −0.62. As should be expected similar results were obtained for the reverse scenario when BayesA was trained in NWL and predictions were made in WL1 (r= −0.60, Table 1). Negative prediction accuracies were not observed for CGM-WGP *(r ≥* 0.22, Table 1), indicating the algorithm was able to define genetic models for physiological adaptive traits that account for the presence of G×E interactions and the effect of the variable environments on the prediction of yield performance.

Prediction accuracy for both CGM-WGP and BayesA increased with increasing number of estimation environments (Table 1, Table 2). The pattern observed for estimation and prediction using single environment data remains evident when more than one environment is included in the estimation set (Table 2). Negative prediction accuracy was still estimated for BayesA when estimation and prediction environments were dissimilar but not for CGM-WGP (Table 2).

### 3.3 Prediction accuracy depends on environment type that reveals genetic variation in adaptive physiological traits

Prediction accuracy for physiological traits depends on the environment type of the estimation set (Table 3). While highest accuracies for BKP were estimated when estimation sets were drought environments WL1 and WL2, highest accuracy for AMAX and RUE were estimated when the estimation set was NWL environment (Table 3). Yield increased markedly with increasing AMAX and RUE within the NWL environment (Fig. 3), which is associated with the highest accuracy attained for these two traits in this environment type. Because the magnitude of the yield response to change in MEB depends on intensity of water deficit at flowering time (Fig. 3, Fig. 2), the highest accuracy for the estimation of MEB was observed when data from WL1, which experienced the greatest level of water deficit at flowering, was used as the estimation set (Fig. 2; Table 3). With the absence of (NWL) or moderate (WL2) water deficit at flowering time, low prediction accuracy was estimated for MEB when these environments were used as estimation sets, −0.03 and 0.17, respectively. These differential accuracies for estimating physiological traits translate into variable accuracies for yield prediction. While accuracy of yield prediction for environment WL2 using a model trained using data for WL1 is 0.51, the accuracy for the reciprocal is 0.34 (Table 2). Similarly, the accuracy of prediction in WL1 using a model trained using data from WL2 and NWL is 0.72, which is lower than for other combinations (0.79 and 0.76). Overall accuracy for yield and physiological trait prediction increased with increasing number of environments included in the estimation set (Tables 1, 2, 3, 4) as more environment types are included that expose variation in adaptive physiological traits enabling the algorithm to find adequate genetic models. This new evidence suggests that further improvements in predictability may be possible by leveraging managed stress environments and optimizing the combinations of types of environments required to expose genetic variation for adaptive traits.

**Table 3.** Mean prediction accuracies (± standard deviation) estimated by the correlation coefficient *r* for physiological process parameters: size of the largest leaf within the leaf area profile (AMAX), mass of ear at silking (MEB), radiation use efficiency (RUE) and limited transpiration trait breakpoint (BKP).

| Estimation<br>environment | Physiological process parameter |  |  |  |
| --- | --- | --- | --- | --- |
|  | AMAX | MEB | RUE | BKP |
|  | ----cm<sup>-2</sup>---- | ----g---- | ----g MJ<sup>-1</sup>---- | ----kPa---- |
| WL1 (1988) | 0.31±0.16 | 0.52±0.15 | 0.33±0.16 | 0.65±0.14 |
| WL2 (2012) | 0.26±0.2 | 0.17±0.2 | 0.33±0.29 | 0.68±0.23 |
| NWL (2010) | 0.51±0.13 | -0.03±0.19 | 0.66±0.13 | 0.17±0.21 |

**Table 4.** Mean prediction accuracies (± standard deviation) estimated by the correlation coefficient *r* for physiological process parameters: size of the largest leaf within the leaf area profile (AMAX), mass of ear at silking (MEB), radiation use efficiency (RUE) and limited transpiration trait breakpoint (BKP).

| Estimation<br>environment | Physiological process parameter |  |  |  |
| --- | --- | --- | --- | --- |
|  | AMAX | MEB | RUE | BKP |
|  | ----cm<sup>-2</sup>---- | ----g---- | ----g MJ<sup>-1</sup>---- | ----kPa---- |
| WL1 + NWL<br>(1988 & 2010) | 0.43±0.14 | 0.53±0.12 | 0.65±0.09 | 0.58±0.12 |
| WL1 + WL2<br>(1988 & 2012) | 0.42±0.10 | 0.46±0.14 | 0.51±0.11 | 0.78±0.07 |
| WL2 + NWL<br>(2012 & 2010) | 0.47±0.11 | 0.28±0.18 | 0.66±0.10 | 0.77±0.07 |
| WL1 + WL2 +<br>NWL (1988 &<br>2010 & 2012) | 0.44±0.12 | 0.51±0.12 | 0.60±0.08 | 0.75±0.07 |

### 3.4 CGM-WGP improved prediction accuracy relative to BayesA for systems where G×E is an important determinant of yield is demonstrated in breeding trials

As in the simulation study above, for the empirical study prior to analysis of the grain yield data and application of the CGM-WGP and BayesA methods the impact of the different irrigation management strategies was investigated to characterize the two environments. Following the same procedures described above for the simulation study, the daily time step water SD profiles were determined for the Viluco well-watered and water-limited irrigation treatments (Fig. 4). For the well-watered treatment the SD profile remained at, or close to, 1.0 for the duration of the crop cycle. Thus, the well-watered treatment was managed to realize a nonwater-limited environment (NWL). For the water-limited treatment irrigation was reduced, commencing in the vegetative stage around V7, and the SD profile decreased to a value below 0.2, coinciding with the timing of flowering for the DH populations tested. After flowering of the DH families was completed irrigation was resumed and the SD profile increased and was maintained between 0.5 and 1.0 for the remainder of the crop cycle. Thus, the water-limited treatment was managed to realize a water-limited environment (WL), where the peak of the water limitation was imposed during flowering time of the DH lines to impact yield development through imposition of a flowering stress drought. Therefore, the testing of the four DH populations in the combination of the NWL and WL environments provided a suitable empirical MET for evaluating the CGM-WGP methodology, where the traits of interest were relevant for predicting grain yield of the maize DH lines in drought and non-drought environments, as was demonstrated in the simulation study.

**Figure 4.**
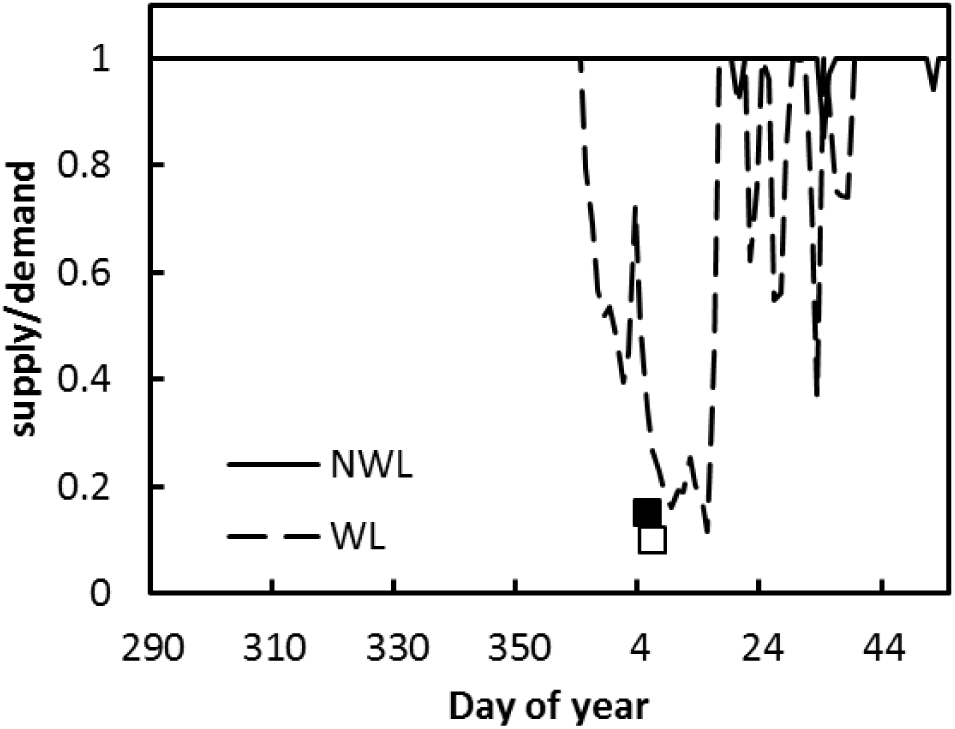
Water supply to demand ratios for DuPont Pioneer Viluco research station in 2012 calculated for water limited (WL) and not water limited (NWL) conditions using a crop growth model as a function of day of year. Simulated shedding date for NWL and WL treatments indicated as closed and open squares. Model parameters were set to means of values utilized in sensitivity analyses, size of the largest leaf in the leaf area profile (AMAX)=950, ear size at silking (MEB)=0.8, radiation use efficiency (RUE)=1.8, and limited transpiration breakpoint (BKP)=2.25.

Analysis of variance indicated that there were significant G×E interactions for grain yield between the NWL and WL environments for all four DH populations. Therefore, further analyses of variance for each DH population focused on the genetic correlation for yield between the two environments and the magnitude of genetic variance for yield within the NWL and the WL environments (Table 5). There was significant genetic variation for grain yield in the NWL and WL environments for all four DH populations. The magnitude of the variance components for yield in the WL environment was greater for DHPop3 and DHPop4 compared to DHPop1 and DHPop2 (Table 5). For the NWL environment, the magnitude of genetic variance for yield was greater for DHPop4 than for the other three DH populations. The genetic correlation for grain yield between the NWL and WL environments differed among the four DH populations (Table 5). To visualize the different magnitudes of genetic variance for grain yield and the different genetic correlations between the NWL and WL environments scatter diagrams were constructed for each DH population for the grain yield BLUPs (Fig. 5). For DHPop1 (Table 5, Fig. 5a) there was no correlation between NWL and WL. For DHPop2 there was an indication of a negative genetic correlation (Table 5, Fig. 5b). For both DHPop3 (Table 5, Fig. 5c) and DHPop4 (Table 5, Fig. 5d) there was a positive correlation. The different grain yield results for the four DH populations provides evidence that the physiological and therefore the genetic basis of the grain yield variation expressed in the NWL and WL environments differed among the four DH populations. Therefore, the results obtained from the empirical MET encompass a number of the scenarios examined in the simulation study and represent important technical issues that need to be addressed in an applied maize breeding program.

**Figure5.**
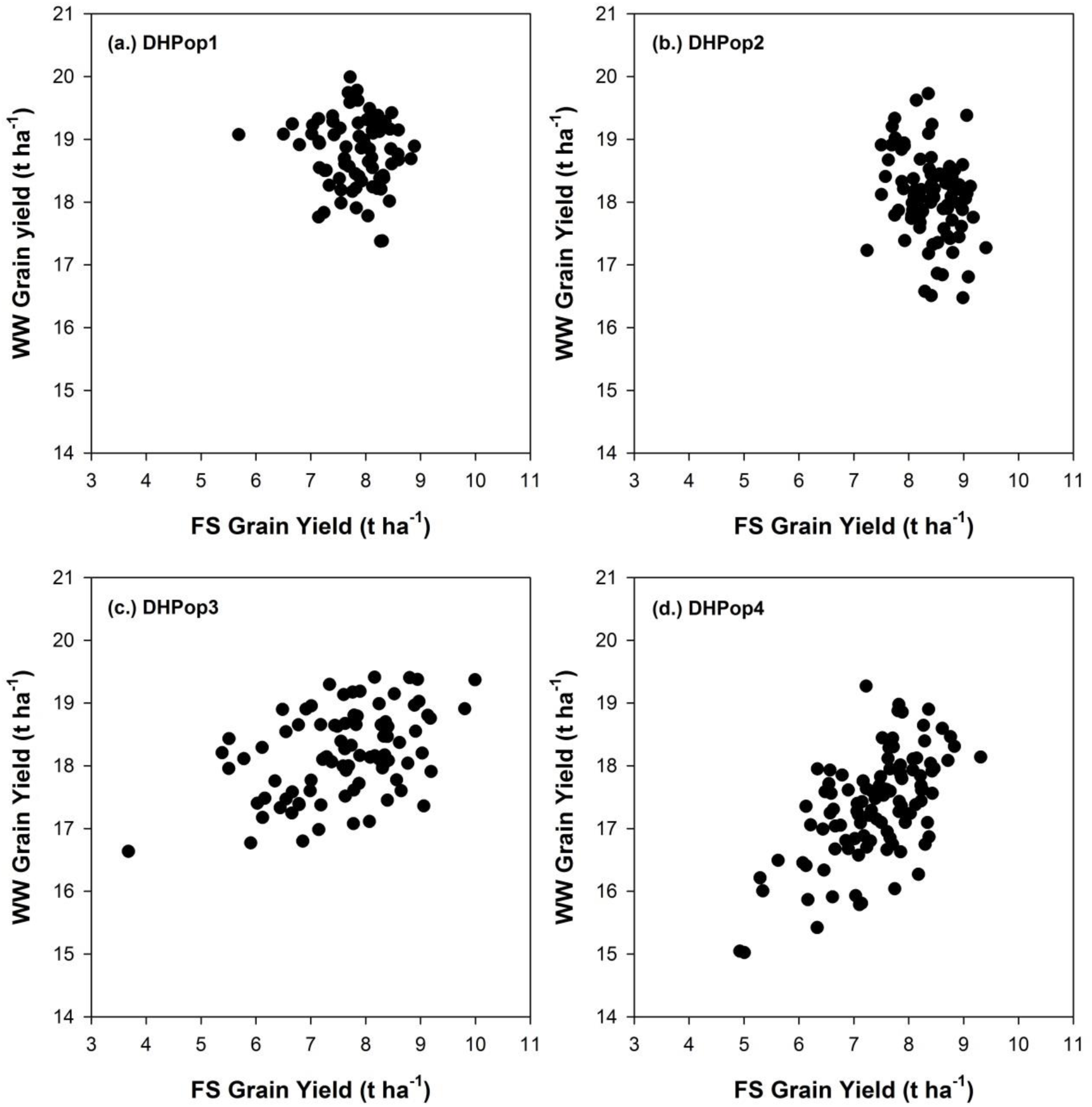
Grain yield observed under water-limited at flowering (WL) and non-water-limited (NWL) due to well watered (WW) environmental conditions for four DH families evaluated at the DuPont Pioneer Viluco research station in 2012.

**Table 5.** Estimates of the genetic correlation between the non-water-limited (NWL) and water-limited (WL) environments (r_G(NWL,WL)_ ± Standard Error), genetic variance component within the non-water-limited environment (VG_NWL_ ± Standard Error) and genetic variance component within the water limited environment (VG_WL_ ± Standard Error) for grain yield (t ha^−1^) of four DH populations tested under WL and NWL environments at the DuPont Pioneer Viluco research station in 2012.

| DH Population | $r_{G(NWL,WL)}$ | $VG_{NWL}$ | $VG_{WL}$ |
| --- | --- | --- | --- |
| DHPop1 | $-0.08 \pm 0.19$ | $0.515 \pm 0.143$ | $0.505 \pm 0.130$ |
| DHPop2 | $-0.20 \pm 0.17$ | $0.686 \pm 0.157$ | $0.375 \pm 0.103$ |
| DHPop3 | $0.36 \pm 0.13$ | $0.689 \pm 0.161$ | $1.433 \pm 0.264$ |
| DHPop4 | $0.49 \pm 0.11$ | $0.944 \pm 0.179$ | $0.920 \pm 0.168$ |

On average CGM-WGP (*r* = 0.34) resulted in higher prediction accuracy than BayesA (r = 0.23) over all predication scenarios considered for the four DH populations (Table 6). Both CGM-WGP and BayesA had higher prediction accuracy for the WL environment (*r* = 0.45 and 0.33, respectively) than for the NWL environment (*r* = 0.23 and 0.14, respectively). In the WL environment the prediction accuracy was positive for all DH populations for both CGM-WGP and BayesA (Table 6). Further for all four DH populations the prediction accuracy for the CGM-WGP was higher than for BayesA for the WL environment. For the NWL environment CGM-WGP achieved positive prediction accuracy for all four DH populations. However, for BayesA the ability to achieve a positive prediction accuracy for the NWL environment depended on the DH population. For two of the DH populations (DHPop1 and DHPop3) positive prediction accuracy was achieved, while for the other two DH populations (DHPop2 and DHPop4) it was not possible to predict grain yield in the NWL environment. Collectively the prediction accuracy results indicate that compared to the BayesA methodology, which was applied to the average grain yield performance across the WL and NWL environments, there were realized advantages in prediction accuracy for yield in both the WL and the NWL environments from the modeling of the G×E interactions between the WL and NWL environments by the CGM-WGP methodology.

**Table 6.** Average prediction accuracy for the CGM-WGP and BayesA methodologies obtained from applying the leave-one-family-out cross-validation approach to prediction of grain yield within the non-water-limited (NWL) and water-limited (WL) environments for four DH families.

| DH family | CGM-WGP |  | BayesA |  |
| --- | --- | --- | --- | --- |
|  | WL | NWL | WL | NWL |
| DHPop1 | 0.28 | 0.16 | 0.16 | 0.30 |
| DHPop2 | 0.44 | 0.38 | 0.29 | -0.06 |
| DHPop3 | 0.64 | 0.21 | 0.50 | 0.29 |
| DHPop4 | 0.45 | 0.18 | 0.36 | 0.02 |

### 3.5 Evaluation of germplasm in virtual environments augments empirical testing through simulation of untested hybrids in large scale evaluation trials

In this paper a set of yield predictions were made to compare CGM-WGP methodology with BayesA using a breeding population (Table 6). Because yield predictions were made utilizing the CGM-WGP methodology, parameters to run the mechanistic CGM could be estimated for these lines from genotypic information and allele values. Figure 6a shows simulated yields for two DH lines in 2263 grids in 2014 and 1988. Yield for DH line 1 showed higher yields than DH line 2 virtually across the U.S. corn belt in 2014 but not in 1988. The yield differential varied with geography indicating G×E interactions. Indeed, the relationship between DH line 1(*x*) and 2 (*y*) across all grids and the two years characterized using a linear regression model is *y* = 51.8(+̅0.6) + 0.94(+̅0.00051)*x*. From these parameters it is feasible to estimate that DH line 1 perform better than DH line 2 in environments where yield is greater than 877 g m^−2^ but not below this threshold. Further evaluations were conducted for two years and 34 DH lines (Fig. 6). This level of testing is not feasible using empirical approaches even at advanced stages of product evaluation, but views of plausible performance that can inform testing could become available for untested DH lines. These results demonstrate reduction to practice of the methodology at the scale of the early hybrid advancement stages of a breeding program. The method demonstrated in this paper enable simulation studies that inform decision about the creation of products and their plausible placement in different geographies and cropping systems.

**Figure 6.**
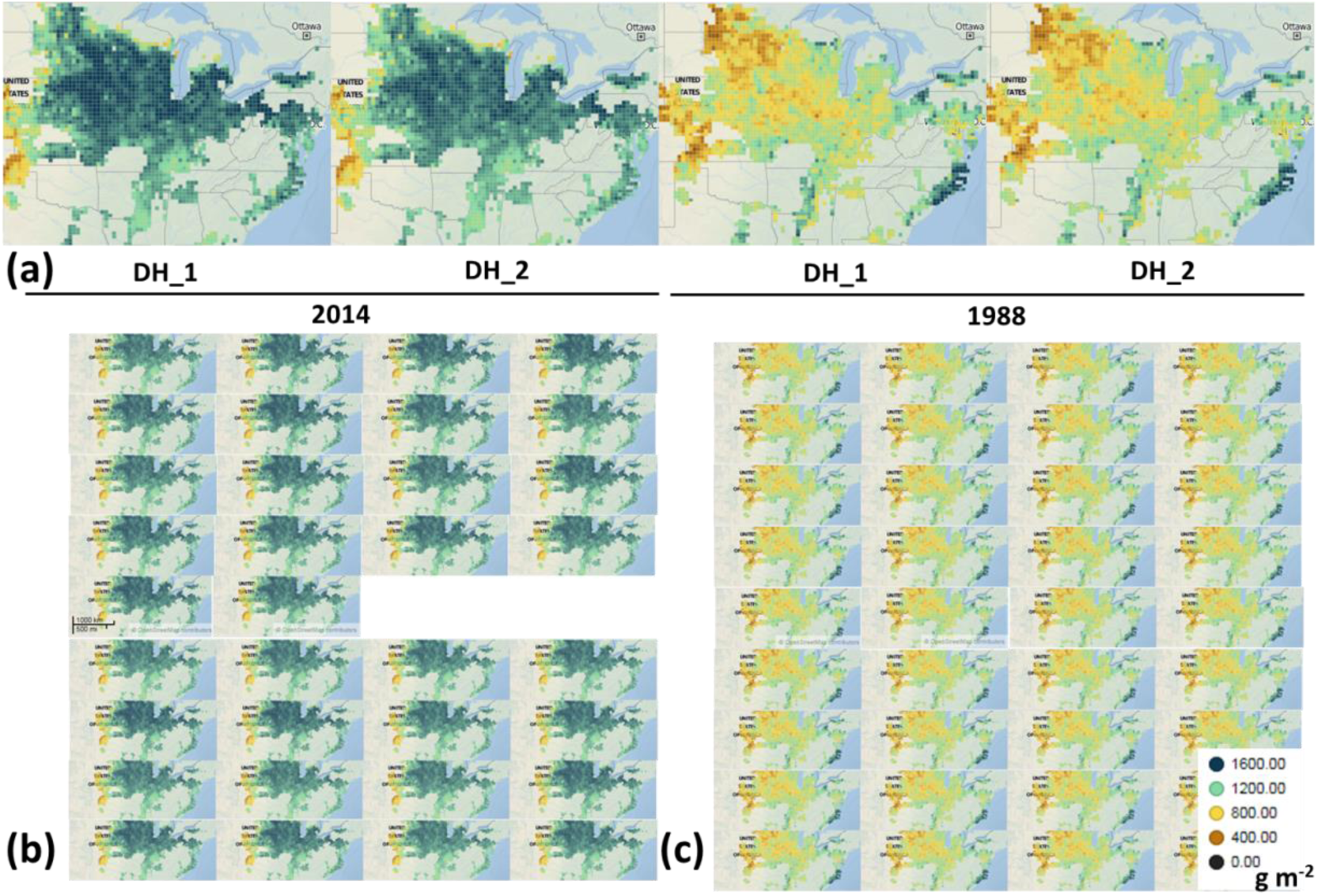
Simulated yields for the major maize growing regions in the U.S. corn belt for 2014 (a,b) and 1988 (a,c) and individuals of a breeding population with CGM parameters estimated from maker data using the CGM-WGP methodology.

## 4. Implication for crop model development and application

The integration of the hierarchical model with the Metropolis-Hastings within Gibbs algorithm enabled using multiple sources of data, observed and virtual, to evaluate CGM-WGP prediction skill. This method is a significant advancement relative to and advocated by prior studies (Technow et al., 2015; Cooper et al., 2016). This enhanced capability enabled studies that demonstrated that increasing the number of environments per se does not necessarily increase predictive accuracy of CGM-WGP but the combination of environment types that enable expression of relevant genetic variation for adaptive physiological traits is a productive approach to increase predictive accuracy.

To realize potentially higher predictive skills, CGMs must incorporate quantitative biological relations that can capture the expression of traits when these are exposed as a result of new experimental designs and precision phenotyping. Many of these relations are unknown and likely surface to the known with the creation of new germplasm and genetic diversity. While the construction of the CGM used in this study was guided by prior research on relevant traits influencing yield in WL and NWL environments (Hammer et al., 2009; Messina et al., 2009; Messina et al., 2011; Choudhary et al., 2014; Cooper et al., 2014; Messina et al., 2015; Reyes et al., 2015; Cooper et al., 2016) it was possible to identify variation in predictive accuracy among breeding populations and examples where CGM-WGP was not better than the reference BayesA method. More cases such as this should be expected and suggest the need to consider plausible avenues of research and development to maximize the opportunities to realize improved accuracy. Out of many plausible solutions the development of a “second generation crop growth model” (SCGM) and the link between Phenomics and CGM are considered briefly.

A SCGM for use within CGM-WGP is designed and created within a dynamic framework that enables rapid changes in the CGM to align it to evolving WGP algorithms, germplasm, breeders’ questions and objectives, to leverage phenomics capabilities, and to effectively deal with scientist bias. Brown et al. (2014) demonstrates a promising framework to enable the development of models of various complexities and that could be evolved quickly by practitioners. A missing component to Brown’s framework is a life cycle management system that provides guidance to the needs of creating new models as well as to retire models. The implementation of authoritative repositories of quality experimental data is necessary to a SCGM life management system and eliminates the need for centralized controlled systems that may delay the implementation of SCGM. Holsworth et al. (2014) discuss promising opportunities such as “The Stack” that can enable model evaluation but also simulation at global scale, extending the application demonstrated in this research for the U.S. corrn belt to other crops and geographies.

A SCGM will be applied to experiments with increasing size, data type diversity generated by phenotyping platforms, and complexity. Although the metropolis-within Gibbs sampling algorithm introduced here can improve the efficiency and decrease computing demands, as demonstrated in this paper, it is inevitable that faster models will be required to work with larger and more complex breeding experiments, involving hundreds of environments and thousands of genotypes from diverse germplasm sources, where convergence due to a more complex system may require increasing the number of iterations to reach convergence. Runtime will become even more relevant as stochasticity is incorporated into SCGM to deal formally with measurement error, environmental uncertainty, and internal stochasticity characteristic of complex systems (Wallach et al., 2012; Technow et al., 2015). Such models in combination with faster algorithms will enable answering fundamental questions about the role of internal system variability on the determination of predictability. Both the need to explore larger CGM parameter spaces when applying the model to new breeding populations, and the availability of trait information that will force identification of solutions that satisfy multiple constraints suggests that a fundamental area of research can focus on how predictability and prediction skill changes with CGM-WGP complexity. As model complexity increases, so will the unknowns and the complexity of the performance landscape that may reduce the ability to find global optima. Answers to these questions will enable designing improved strategies to effectively deal with limited predictability to accelerate genetic gain.

Phenomics and SCGM will be integrated to form a virtuous cycle of mutual development while delivering improved genetic and physiological mechanistic models, and improved predictions (Cooper et al., 2002; Houle et al., 2010; Messina et al., 2011). High throughput phenotyping platforms, here considered broadly to include managed stress environments, will provide data to both train CGM-WGP and to enhance biological mechanistic models (Houle et al., 2010; Cooper et al., 2014; Pauli et al., 2016). CGM can inform phenotyping based on known physiological understanding but also serve as frameworks that help identify knowledge gaps (Hammer et al., 2002). SCGM-WGP becomes the integration framework that enables the incorporation of data for multiple traits, which are outputs from high throughput phenotyping platforms, genomic information, agronomy and environment. A model to enable such integration could be of the form,

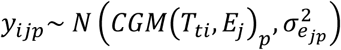

where *y_ijp_* is a set of measurable phenotypes (*p*) influenced by functional trait *t* and measurable using platforms (eg. leaf area, leaf angle that influence functional traits, such as RUE, which are captured in the relationships of the CGM) with error variance 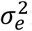 for phenotype *p* in environment *j*. The methods described in this paper could be applied to train this model. This approach is a significant change relative to the two stage approaches advocated to date (Yin et al., 2004; Hammer et al., 2006; Chenu et al., 2009; Messina et al., 2011; Pauli et al., 2016) because it provides an integrated and flexible framework to tune the utilization of phenomics to the breeding or more general, to the plant science objective. Predictions from this model are testable hypothesis with physiological and genetic components. It is less clear in the literature and a clear area of future research how to extend the framework to handle knowledge gaps through integration of polygenic terms that could be handled through statistical models within the framework.

## 5. Concluding remarks

This research used a combination of simulation and empirical studies to explain how the CGM-WGP methodology introduced by Technow et al. (2015) works to achieve enhanced levels of prediction accuracy and to demonstrate reduction to practice for some applications to maize breeding and product placement for target environments in the US corn-belt. This new development in modeling, achieved through the integration of genetic and physiological modeling capabilities, provides a research platform for enhancing predictions of crop yield that can account for important components of G×E×M interactions that influence crop productivity. The development of the hierarchical model enabled studies to demonstrate the critical use of diverse environments as estimation sets for CGM-WGP as these reveal genetic variation for adaptive physiological traits. Three important outcomes of the CGM-WGP methodology that merit further consideration and investigation are: (1) The algorithm used within the CGM-WGP method to connect genetic information to the traits that vary among genotypes and respond to environmental conditions to influence yield is conducted as an integrated single step process. This single step approach contrasts with previous two-step methods that first seek to identify genes or regions of the genome (QTL) that map to traits proposed to influence yield, which are then in turn integrated within the CGM as predictors in a second step. (2) Some of the requirements of high-throughput phenotyping are relaxed and radically changed in comparison to the alternative two-step approaches. In the two-step approaches advocated to date all individuals have to be phenotyped for all traits in all environments to establish the relationship between genes or QTL and the traits prior to integration into the CGM. For CGM-WGP this dense level of phenotyping may not always be required, the phenotyping load can be reduced, and efforts seeking to generate large volumes of data may be turned into efforts seeking to unravel the physiological basis of adaptation and how to incorporate the resulting knowledge into mechanistic models. For many applications emphasis can be placed on phenotyping efforts to establish the functional relationships that are established within the CGM and to develop informative prior distributions to be used with the CGM to enable the CGM-WGP. (3) New criteria are emerging as foundational principles for developing a second generation of CGMs that explicitly incorporate the concept of genetic variation for traits at different scales in biological hierarchies (e.g., cellular to tissue to organ to crop canopy), and their functional relationships across these hierarchical scales in biology.

